# Implicit counterfactual effect in partial feedback reinforcement learning: behavioral and modeling approach

**DOI:** 10.1101/2020.09.30.320135

**Authors:** Zahra Barakchian, Abdol-hossein Vahabie, Majid Nili Ahmadabadi

## Abstract

Context by distorting values of options with respect to the distribution of available alternatives, remarkably affects learning behavior. Providing an explicit counterfactual component, outcome of unchosen option alongside with the chosen one (Complete feedback), would increase the contextual effect by inducing comparison-based strategy during learning. But It is not clear in the conditions where the context consists only of the juxtaposition of a series of options, and there is no such explicit counterfactual component (Partial feedback), whether and how the relativity will be emerged. Here for investigating whether and how implicit and explicit counterfactual components can affect reinforcement learning, we used two Partial and Complete feedback paradigms, in which options were associated with some reward distributions. Our modeling analysis illustrates that the model which uses the outcome of chosen option for updating values of both chosen and unchosen options, which is in line with diffusive function of dopamine on the striatum, can better account for the behavioral data. We also observed that size of this bias depends on the involved systems in the brain, such that this effect is larger in the transfer phase where subcortical systems are more involved, and is smaller in the deliberative value estimation phase where cortical system is more needed. Furthermore, our data shows that contextual effect is not only limited to probabilistic reward but also it extends to reward with amplitude. These results show that by extending counterfactual concept, we can better account for why there is contextual effect in a condition where there is no extra information of unchosen outcome.

## Introduction

In everyday life, we frequently decide between options. Value of an option is usually learned via trial and error [1], and it is represented in multiple cortical and subcortical areas of the brain. Values of competing options interact with each other and consequently the context in which options are located can affect the representations [2]. Although the early studies on the contextual effects have been designed in the decision-making paradigm [3–8], a new trend has been formed recently to show that some of the behavioral biases come from contextual effects during value learning [9–11]. They showed that, in particular, in the paradigm in which the counterfactual outcomes pertaining to the chosen option were available (Complete feedback), subjects were strongly affected by the context, and this is mostly because they were induced to use a relative strategy. Although, it has been shown that there is a weaker contextual effect in the Partial version [9, 11], yet there is no global consensus about whether and how this effect happens.

Reinforcement learning is an incremental procedure that updates the option value via its prediction error [1]. This procedure is happening in the striatum where encodes action values [12–17], and is modulated by dopamine which encodes prediction errors [18]. Dopamine has opposing exciting and inhibiting effects on two distinct populations of striatal neurons called D1-SPNs and D2-SPNs (spiny projection neurons) respectively [19, 20]. Some reinforcement learning studies have shown opposing activities with similar strength in these two clusters during learning [21, 22]. Recent optical evidence suggests a model for Basal Ganglia that, it is the relative activity of these two clusters that represents an internal decision variable during decision making and learning [23–25]. For a good review on this issue, see [26]. Inspired by the opposing role of dopamine on D1- and D2-SPNs, while they encode two competing options’ values, we proposed a simple reinforcement learning model called Opposing Learning model, in which the chosen prediction error in addition to updating the chosen option value (classic standard Q-learning), updates the unchosen option value, though in an opposing manner. This mechanism is consistent with diffusive nature of dopamine release and enables the model to endogenously encode the chosen and unchosen options’ values relative to each other and consequently suggests having a contextual effect in the Partial feedback conditions too.

In the Complete feedback paradigm in which there are some exogenous factors that impose relativity on value learning procedure, the value learning strategy can be complex [9–11]. In these conditions, the main strategy might be to compare two presented outcomes and this would generate the regret and relief emotions. It has been shown that people tend to optimize their outcome difference, outcome_factual_ - outcome_counterfactual_ (i.e. minimize their regret and maximize their relief) [27–29]. In the Partial feedback paradigm, due to absence of regret and relief emotions, the main value learning strategy assumes to be the standard maximizing expected rewards. Interestingly, recent studies have illustrated that people are neither fully expected-reward optimizer nor fully outcome-difference optimizer, rather they are hybrid optimizers [9, 30], who use both of these strategies with different weights. The individual differences among people would depend on how much a person weighs each of these strategies. By adding a hybrid component to the basic OL model, we could extend the OL model for the Complete version too.

In this paper, we went beyond the standard definition of a counterfactual outcome, and focused on an uncommon subtle aspect of counterfactual role in value learning. This role is important in particular in the situations where there is no forgone outcome to trigger the comparison-based strategy explicitly. We used two types of feedback paradigms, with and without forgone outcomes. By using the chosen outcome as a counterfactual outcome for unchosen option we introduced a novel reinforcement learning model that could account for the contextual effect of the behavioral data better than previous related models. To see how contextual effect is different in two types of cortical and subcortical dominant behavior, we evaluated participants’ behavior in two post-learning transfer phase and value estimation phase. We observed that participants behaved strongly biased in the transfer phase, while this bias was very weak in the estimation phase. This suggests that these two systems have different sensitivities to the contextual effect, such that subcortical system is more sensitive than cortical one. To better dissociate the cortical and subcortical behavioral difference, we used reward amplitude rather the reward probability, because we assumed that complexity of reward amplitude can better engage the cortical parts of the brain in the estimation phase. Thus we could show that there is contextual effect also for reward amplitude.

## Results

### 1 Results

#### Behavioral task

Two groups of participants performed two different versions of the instrumental learning tasks, the Partial feedback version, in which the factual outcomes were only provided, and the Complete feedback version, in which both factual and counterfactual outcomes were provided. Subjects were instructed to gain the most possible rewards during the task. Rewards were random independent numbers drawn from particular normal distributions. They observed two pairs of options (*A*_1_, *B*) and (*A*_2_, *C*), where *A*_1_ and *A*_2_ were associated with rewards from the same distribution of 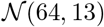 and *B* and *C* were associated with rewards from two different distributions of 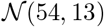 and 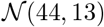 respectively. To conceal the task structure from the participants, although their associated values were equal, the images assigned to *A*_1_ and *A*_2_ were different. After the learning phase, they unexpectedly entered the post-learning transfer phase, in which all possible binary combinations of options (6 pairs) were presented to them (each combination 4 times), and they were asked to choose the option that was associated with the highest expected rewards in the preceding learning phase. With this design, if there is a bias in preferring *A*_2_ over *A*_1_, this transfer phase can reveal it. Similar designs were used in the context-dependent learning studies as well [9–11]. In order not to interfere with their previous learning, no feedbacks were provided in the transfer phase [9–11, 31, 32]. After each choice, they were asked to report their choice confidence on a scaled bar from 0 to 100. Finally, in the value estimation phase we asked the subjects to report their estimated expected value of each stimulus on a scaled bar ranging from 0 to 100.

#### Performance

To see whether the participants learned the options values during the task, at first, we calculated their performance in the learning phase which is the percentage in which they chose the advantageous option (the option with higher expected rewards). We observed that in both versions, the participants’ performances were higher than 0.5. (Partial: performance = 0.7613 ± 0.1130; Complete: performance = 0.8823 ± 0.0853; Figure 3b). Consistent with the previous studies [9–11], we found that in the Complete version, the performance of the learning phase was significantly higher than that of the Partial version (*p* = 4.5603*e* − 07, *tstat* = 5.3522, *df* = 75, one-tailed ttest), which means providing counterfactual outcomes facilitate learning. In addition to the learning phase, we also observed that subjects had higher performance in the transfer phase, such that participants significantly preferred the option with higher expected rewards in each combination (Partial: *p* = 1.0577*e* − 73, *tstat* = 23.4715, *df* = 348; Complete: *p* = 4.3483*e* − 84, *tstat* = 24.7863, *df* = 418; ttest). Additionally, the reported confidences for the most advantageous options were significantly higher than those with non-advantageous options (Partial: *p* = 1.5970*e* − 06, *tstat* = 4.9705, *df* = 173; Complete: *p* = 3.7111*e* − 09, *tstat* = 6.1597, *df* = 208; ttest). For these and all the following analyses, unanswered trials of the learning phase were excluded.

**Fig 1.**
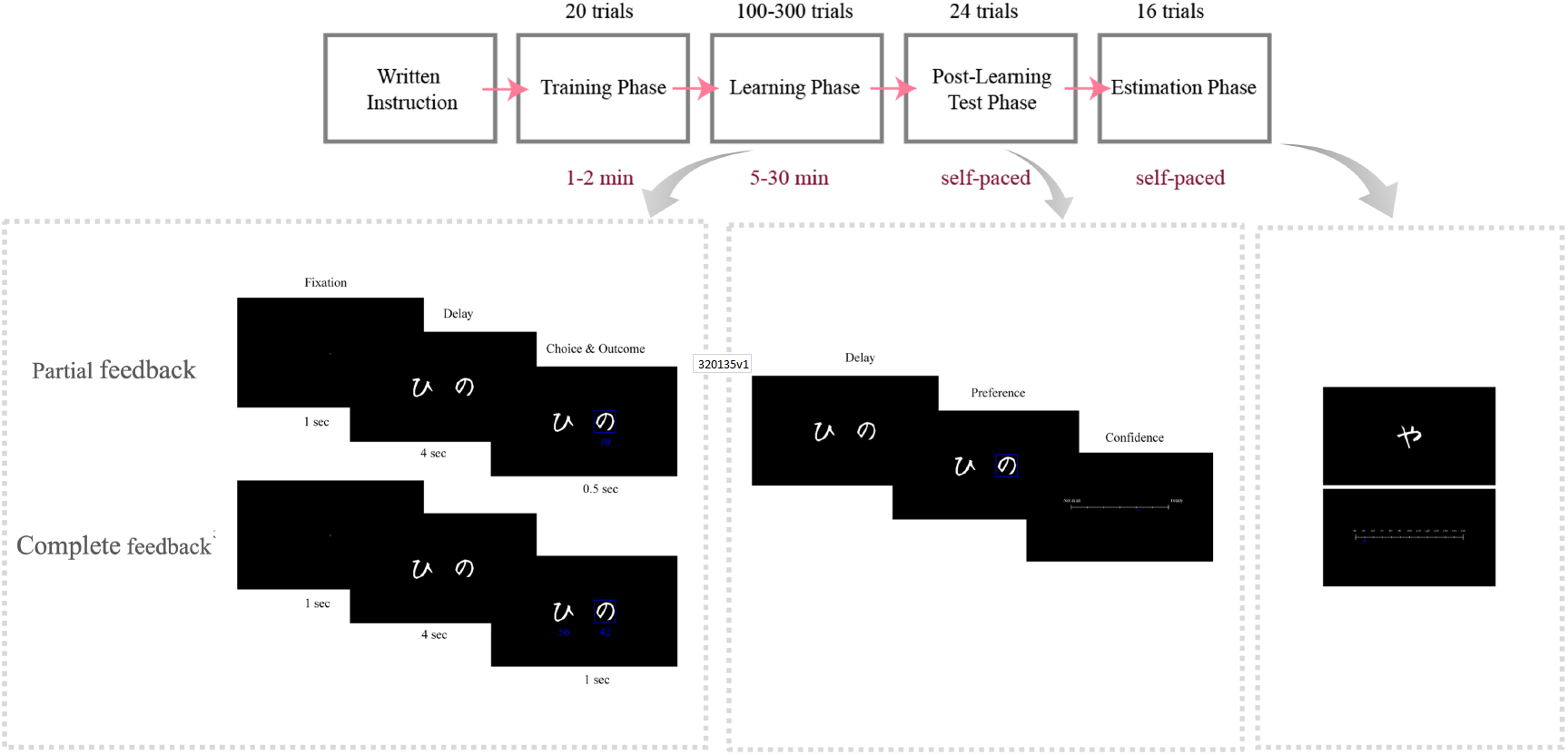
Behavioral Design. Time-lines of the Partial and Complete feedback versions. Subjects were instructed with written instructions and trained through 20 trials before beginning the main task. They learned two pairs of options in the Learning phase with trial and error. They transferred to the transfer phase after at least 100 and at most 300 trials, in which they were supposed to choose the most advantageous option between the two presented options, and report their choice confidence. In this phase, all possible binary combinations of options were presented.

**Fig 2.**
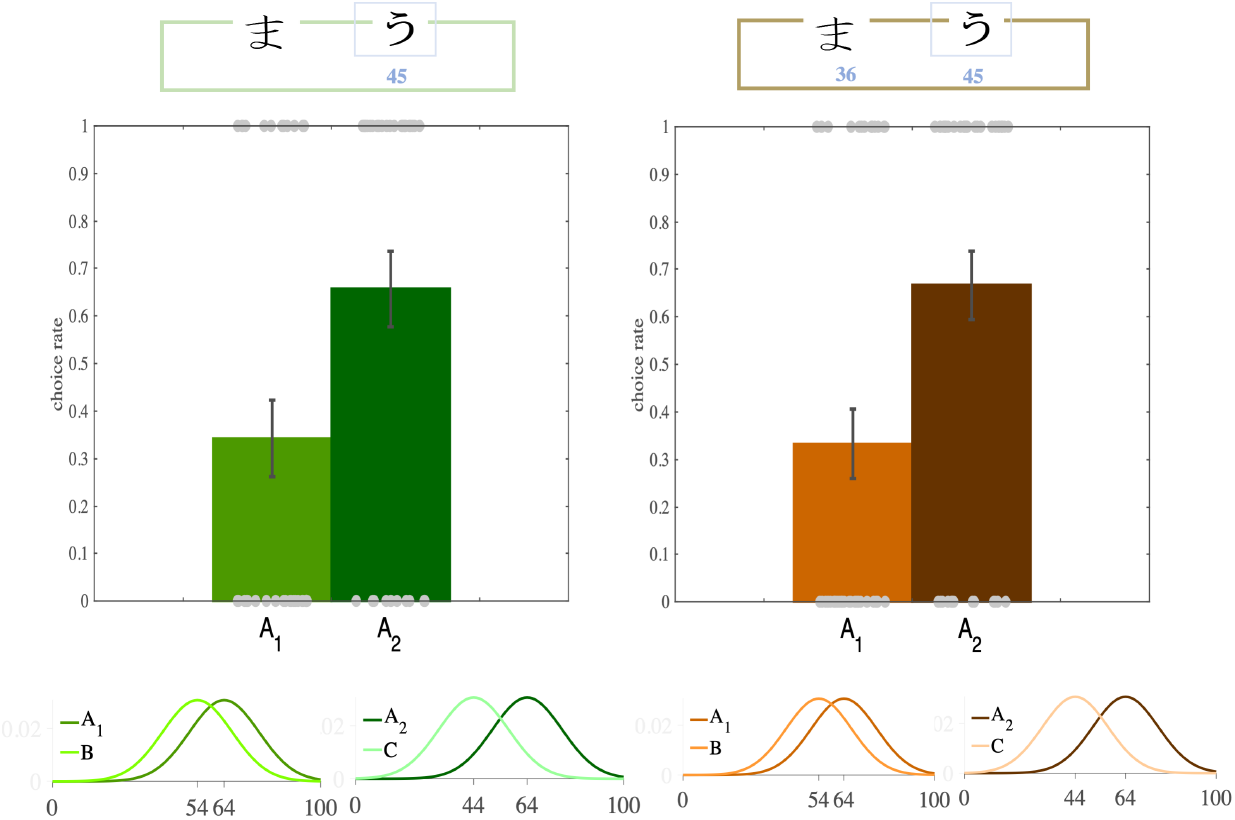
Transfer effect. In the transfer phase of both feedback versions, participants significantly preferred the option with higher relative value (*A*_2_, dark green) between the two options with equal absolute values.

**Fig 3.**
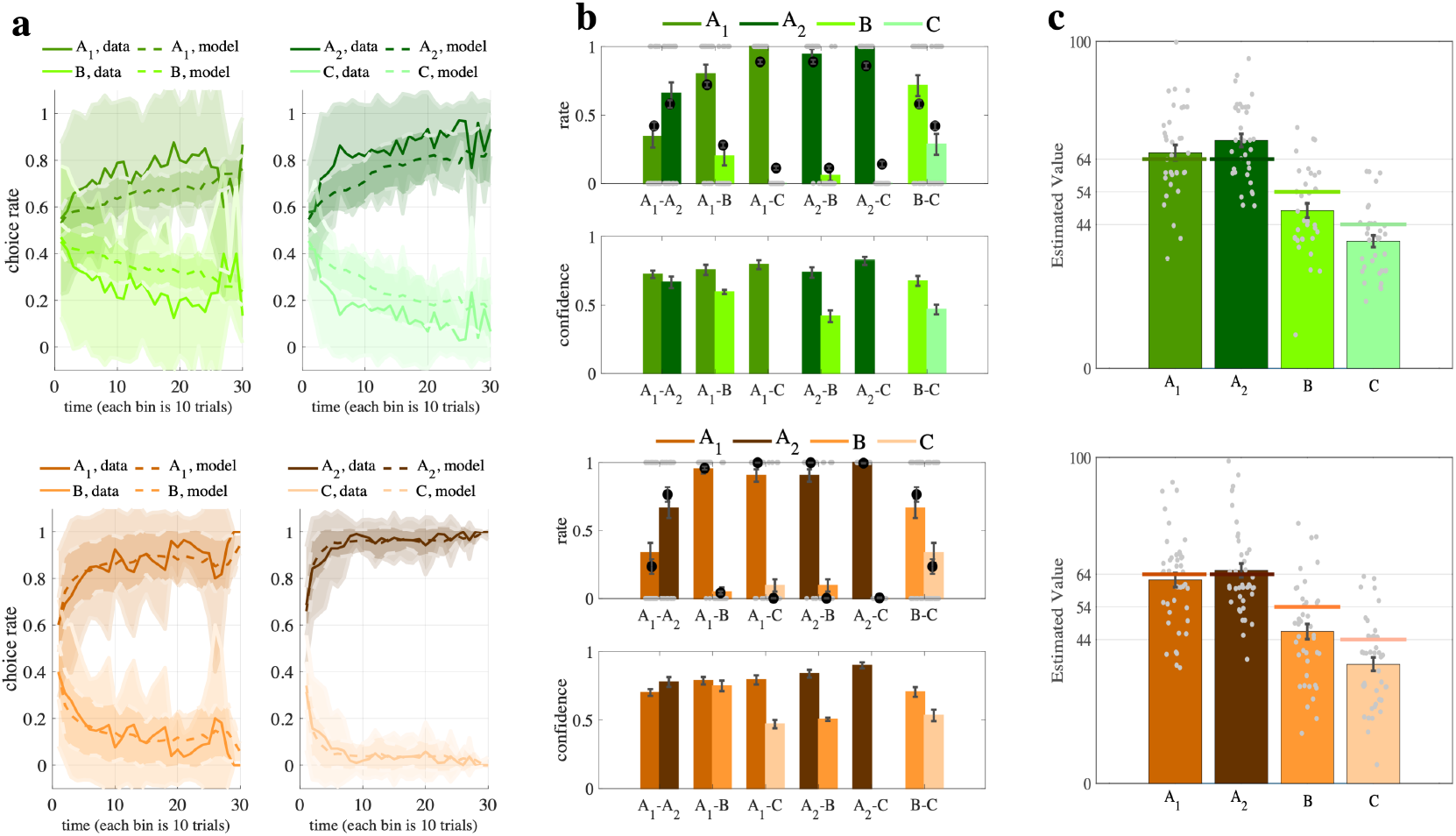
Behavioral results in the learning, transfer and estimation phases. **a.** The learning curves of two pairs of options in the learning phases of both versions show that participants learned to choose the advantageous options (*A*_1_ in *A*_1_*B* pair and *A*_2_ in *A*_2_*C* pair). The learning curve of the OL models in both versions also shows similar results. Each bin in the x-axis is the mean of the choices in 10 trials. The Partial version is green, and the Complete version is brown. Solid lines show the behavioral effect, and dashed lines show the model effect. **b.** The subjects’ preferences in all 6 combinations (top), and corresponding confidences (down), with OL predictions (black dots). **c.** The value estimations of the subjects (colored bars) are very close to the real expected rewards of *A*_1_, and *A*_2_ options (colored lines). Partial version is green and Complete version is brown. Shadings are *SD* and error bars are *SEM*.

#### Contextual effect

When through performance analysis, we made sure that participants learned the options’ values during the learning phase, we turned to the transfer phase to see whether there is any contextual effect. Considering the first iterations of the participants’ choices in the transfer phase, we found that participants’ preferences between *A*_1_ and *A*_2_ have been significantly modulated by their distance from their paired options, such that despite having equal absolute values, participants preferred *A*_2_ over *A*_1_ (*transfer bias*) in both versions (Partial: *p* = 0.04, *ratio* = 0.65; Complete: *p* = 0.01, *ratio* = 0.66; binomial test) (Figure 3a). This trend remained when we consider all the four iterations of *A*_1_ and *A*_2_ though it loses significance (Partial: *p* = 0.08, *ratio* = 0.63; Complete: *p* = 0.053, *ratio* = 0.64; binomial test). This loss of significance might be due to strategy of balanced choice in subjects to reduce the risk for all four choices of no feedback.

To assure that the observed bias in the transfer phase belongs to the context-dependent value learning, and not to some other confounding factors, we probed which other factors could have affected the subjects’ preference toward *A*_2_. The observed bias might have occurred due to the fact that in the learning phase *A*_2_ has been chosen more frequently than *A*_1_. To test this possibility, we ran a logistic regression analysis to see whether the preference of *A*_2_ over *A*_1_ in the first (*A*_1_, *A*_2_) iteration of the transfer phase was due to the difference between the choice frequency of *A*_2_ over *A*_1_ in the learning phase. This analysis showed that the effect of choice frequency of *A*_2_ over *A*_1_ on the transfer bias is not significant for complete version and near significant for partial version (Partial: *p* = 0.054, *tstat* = 1.92; Complete: *p* = 0.12, *tstat* = 1.54). Significant intercept of the regression confirms the transfer effect, even when choice frequency is controlled (Intercept, Partial: *p* = 0.03, *tstat* = 2.15; Complete: *p* = 0.02, *tstat* = 2.20).

Furthermore, we ran another logistic regression analysis to assess whether the different choice frequencies of options in the last trials of the learning phase (last 20 trials) have made the observed bias in the transfer phase. We again found no significant effect of late choice frequency on the transfer bias (Partial: *p* = 0.56, *tstat* = −0.57; Complete: *p* = 0.29, *tstat* = 1.0473) while intercepts remained near significant (Partial: *p* = 0.06, *tstat* = 1.83; Complete: *p* = 0.03, *tstat* = 2.13). The other alternative for transfer bias justification might be the amount of very small or very large rewards (rewards from the top or bottom of the reward distributions) that affected the transfer bias. Again, using logistic regression analysis, we separately tested whether sum of the observed rewards greater than *μ* + 2.5*σ* and less than *μ* − 2.5*σ* (*μ* and *σ* are the mean and standard deviation of the observed rewards, respectively) could affect the transfer bias. We found no significant effect of large and small rewards in either of the versions (large rewards: [Partial: *p* = 0.40, *tstat* = 0.82; Complete: *p* = 0.62, *tstat* = 0.48], Small rewards: [Partial: *p* = 0.54, *tstat* = 0.60; Complete: *p* = 0.47, *tstat* = −0.71]).

#### Value estimation

Considering the participants’ first estimation in the average estimation phase, we found that participants almost precisely estimated the expected rewards of the most advantageous option based on their mean rewards, but the other non-advantageous options have been significantly underestimated (Figure 3c). This illustrates that when an option is chosen frequently, subjects could either track precisely the mean of its rewards or calculate its value at the moment of estimation. This result was also observed when we considered the participants’ total estimation (the average of the four repetitions for each stimuli).

To test whether the observed contextual bias in the transfer phase, would also be observed in the estimation phase, we ran a paired-ttest analysis on the estimated values. There was no significant difference between subjects’ average estimation of *A*_1_ and *A*_2_ in both versions, yet there was a trend in overestimating *A*_2_ compared to *A*_1_ (Partial: *p* = 0.25, *tstats* = −1.14, *df* = 34; Complete: *p* = 0.28, *tstats* = −1.08, *df* = 41; paired-ttest). These results support the dual value-based system hypothesis, in which the subcortical system (BG) is responsible for stimulus-response association (the behavior that dominated in the transfer phase), and the cortical system (Frontal Cortex) is responsible for average reward computation (the behavior that dominated in the estimation phase) [33–37].

#### Comparison effect

When we observe the consequences of our decision, we compare the outcome of our decision with those alternative decisions we could have made. This comparison would trigger feelings of regret and relief if the outcome of our decision is either better or worse than those of other alternatives respectively. People naturally tend to avoid regret (approach the relief), and when one experiences regret (relief), she will likely switch to the other option (stay in the previous choice) [28, 29].

To test whether the outcomes difference of the previous trial of the same condition has influenced the switching behavior of the current trial in the learning phase, we used a hierarchical logistic regression analysis. In this analysis, we modeled the switching behavior of the subjects (1 if subject has switched, and 0 if subject has stayed on her previous choice), as a function of the outcomes difference the subject has experienced in the previous trial of the same condition, and also the difference of the expected values of the options in the current trial. The outcome difference in each trial was defined as the difference of the factual outcome and counterfactual outcome {*r_FC_* − *r_CF_*} in the Complete version, and the difference of the factual outcome and the counterfactual value (expected rewards) {*r_FC_* − *V_CF_*} in the Partial version. All regressors have been z-scored. While this analysis showed that there was a significant comparison effect in the Complete version, it showed no significance in the Partial version (Table 1). This means subjects tend to switch to other alternatives after experiencing regret and stay on their previous choice after experiencing relief, and this tendency is stronger in the Complete version compared to the Partial one. To investigate this effect more thoroughly, we performed a similar analysis on the logarithm of reaction times (logarithm). We observed again that, in the Complete version and not the Partial version, reaction times in each trial were significantly modulated by the outcome difference in the previous trial of the same condition, in a way that whenever the difference is smaller, the reaction time is slower, and vice versa (Table 1). This result is consistent with the post-error slowing phenomena that have been reported frequently in the decision-making literature [38, 39].

**Table 1.** Comparison effect of the subjects’ switch behavior and reaction times.

| *Switch* |  |  |  |  |  |  |  |  |
| --- | --- | --- | --- | --- | --- | --- | --- | --- |
| *partial* |  |  |  |  | *complete* |  |  |  |
| Name | Estimate | SE | tStat | pValue | Estimate | SE | tStat | pValue |
| (Intercept) | -1.5528097 | 0.10551335 | -14.716713 | 2.69E-48 | -2.724902 | 0.17598005 | -15.484153 | 2.28E-53 |
| outcome difference | -0.0879529 | 0.0567055 | -1.5510467 | 1.21E-01 | -0.5462195 | 0.06292942 | -8.6798744 | 4.68E-18 |
| value difference | -1.123403 | 0.08767908 | -12.812668 | 3.67E-37 | -0.9158512 | 0.06505058 | -14.079062 | 1.57E-44 |
| condition | 0.25688705 | 0.08761796 | 2.93189954 | 0.00337999 | 0.25104619 | 0.12809207 | 1.95988856 | 0.05004073 |

| *Reaction Time* |  |  |  |  |  |  |  |  |
| --- | --- | --- | --- | --- | --- | --- | --- | --- |
| *partial* |  |  |  |  | *complete* |  |  |  |
| Name | Estimate | SE | tStat | pValue | Estimate | SE | tStat | pValue |
| (Intercept) | -0.1164283 | 0.03073684 | -3.7879077 | 0.00015321 | -0.1211333 | 0.03585658 | -3.3782727 | 0.00073263 |
| outcome difference | 0.01123051 | 0.00651389 | 1.72408744 | 0.08473669 | -0.0164905 | 0.00526292 | -3.1333433 | 0.00173402 |
| value difference | -0.0699353 | 0.0101347 | -6.9005836 | 5.64E-12 | -0.0698999 | 0.01654412 | -4.2250579 | 2.41E-05 |
| condition | 0.04191482 | 0.02390541 | 1.75336139 | 0.07958424 | 0.03658956 | 0.02364193 | 1.54765513 | 0.12174177 |

The hierarchical logistic regression and hierarchical simple regression analysis were performed on switch behavior and logarithms of reaction times of the subjects respectively. Along with the outcome difference as the main regressor, the current value differences between the two paired options and the condition type ( $A_1B$ , $A_2C$ ) were also included as a control regressor. The results illustrate that both current participants' choices and current reaction times were significantly influenced by the outcome differences of their previous choices in the Complete but not Partial version.

#### Opposing Learning model (OL)

Here, we introduce a novel reinforcement learning model, called **OL model**, adopted from the standard Q-learning model and inspired by the striatal mechanism. At first, we introduce the basic model for the Partial version, and then we extend this model for the Complete version.

##### Model description

This model is chosen-option centered in a way that value updating is done based on the prediction error of chosen option. Following the choice, the chosen prediction error simultaneously updates the chosen and unchosen values in an opposing manner (increasing and decreasing respectively). This mechanism is inspired by the opposing effect of dopamine on D1-SPNs and D2-SPNs neurons in the striatum, where they encode chosen and unchosen options respectively. The main reason to apply a single prediction error for updating of both options is that dopamine release is diffusive and so it is non-selective during release, thus, it will affect both D1 and D2-SPN neurons simultaneously.

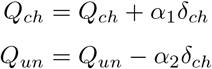

where *ch* referred to *chosen* option, *un* referred to *unchosen* option, and *δ_ch_* = *r_ch_ − Q_ch_*. Generally, when we refer to the OL model, we mean the OL model with two different learning rates, but in this paper, when we want to compare the two different versions of the OL model, *α*_1_ = *α*_2_, and *α*_1_ ≠ *α*_2_, we call them OL_1_, and OL_2_ respectively. When the subject compares the options’ values for making the choice, the decision is made by the softmax rule, 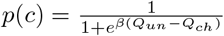 where *β* is the inverse of the temperature parameter. The OL behavior is strongly dependent on the amount of *α*_2_ relative to *α*_1_. Based on *α*_2_ in either of these three intervals: 0 *≤ α*_2_ < *α*_1_, *α*_2_ = *α*_1_, or *α*_1_ < *α*_2_ < 1, the model generates a particular behavior.

##### OL contextual effect

In the OL model, the chosen and unchosen values are coupled, therefore their coding is not independent of each other, rather they are negatively correlated. Our simulation shows that the correlation between two paired options is proportionate to the following formula:

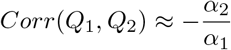

According to this formula, the amount of the correlation between *Q*_1_ and *Q*_2_ depends on the amount of unchosen learning rate. When *α*_2_ changes from 0, where *Q*_1_ and *Q*_2_ are almost orthogonal (*corr* ≈ 0), to *α*_1_, where *Q*_1_ and *Q*_2_ are almost fully correlated (*corr* ≈ −1), the encoding will change from almost fully absolute to almost fully relative (Figure 5a,b). Via simulating the experiment with typical agents of *α*_2_ = 0, 0 < *α*_2_ < *α*_1_, and *α*_2_ = *α*_1_, we showed that we will have zero, moderate and large amount of contextual effect with never, temporary and permanent contextual effect, respectively (Figure 5c).

**Fig 4.**
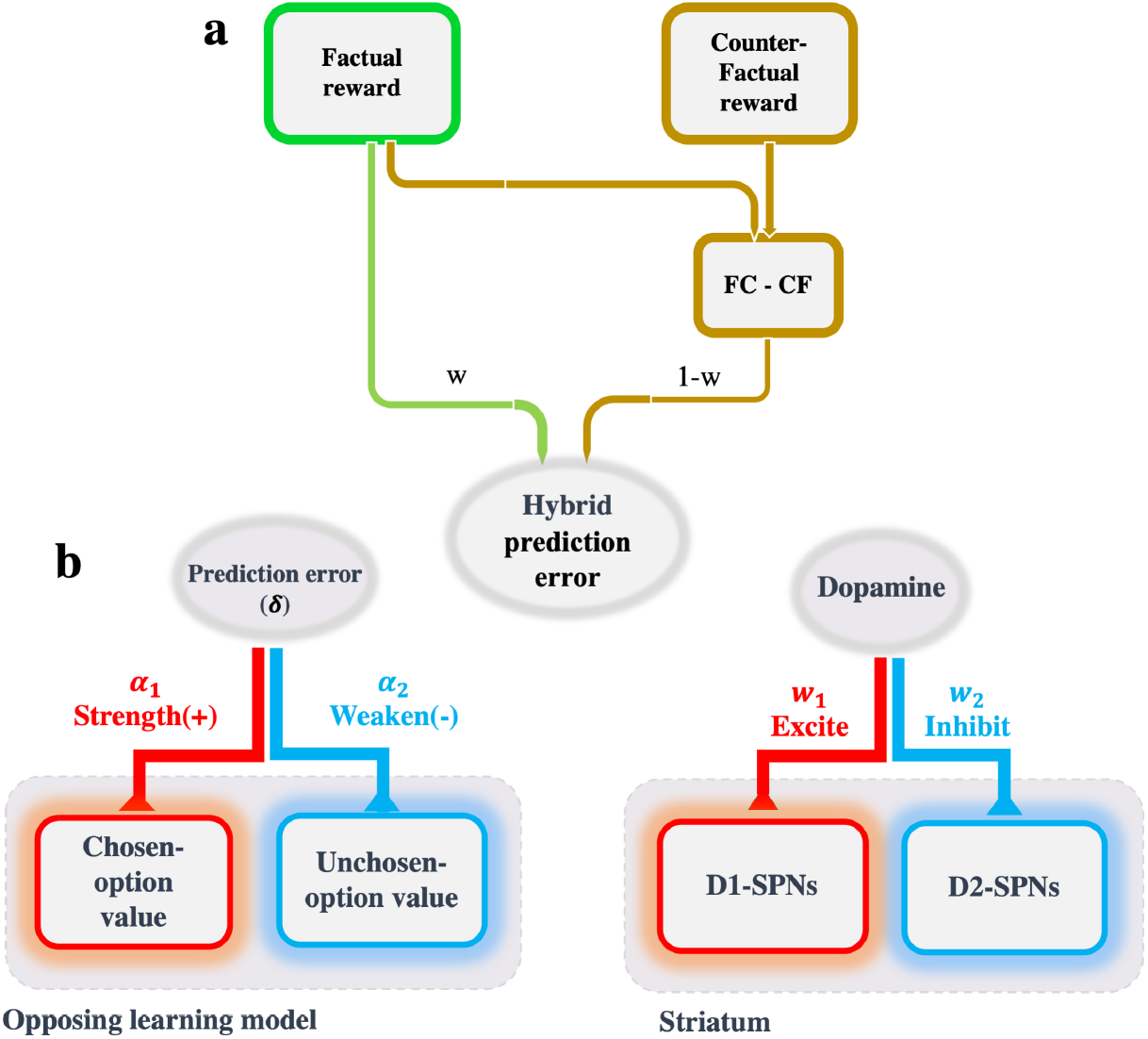
The schematic of the OL model and its extension. **a.** The comparison of the competing outcomes is a common strategy in the value learning strategy, particularly in situations where counterfactual outcomes are also provided along with factual outcomes. This comparison triggers the people’s regret (relief) emotion which subsequently drives the avoidance (approach) action behavior. This tendency to minimize regret (and maximize relief) along with the tendency to maximize the expected rewards as a hybrid strategy that can account for the behavioral data is better than either of these strategies. The weights assigned to each strategy, absolute and relative, determine the amount of its effect on behavior. **b.** The idea behind the OL model comes from the opposing role of dopamine on two distinct populations of D1-SPN and D2-SPN neurons, which have been proposed to encode the chosen and unchosen options’ values, by promoting the latter and inhibiting the former. Correspondingly, in the inspired model, chosen prediction error has an opposing role in updating the chosen and unchosen options’ values, by strengthening the latter and weakening the former.

**Fig 5.**
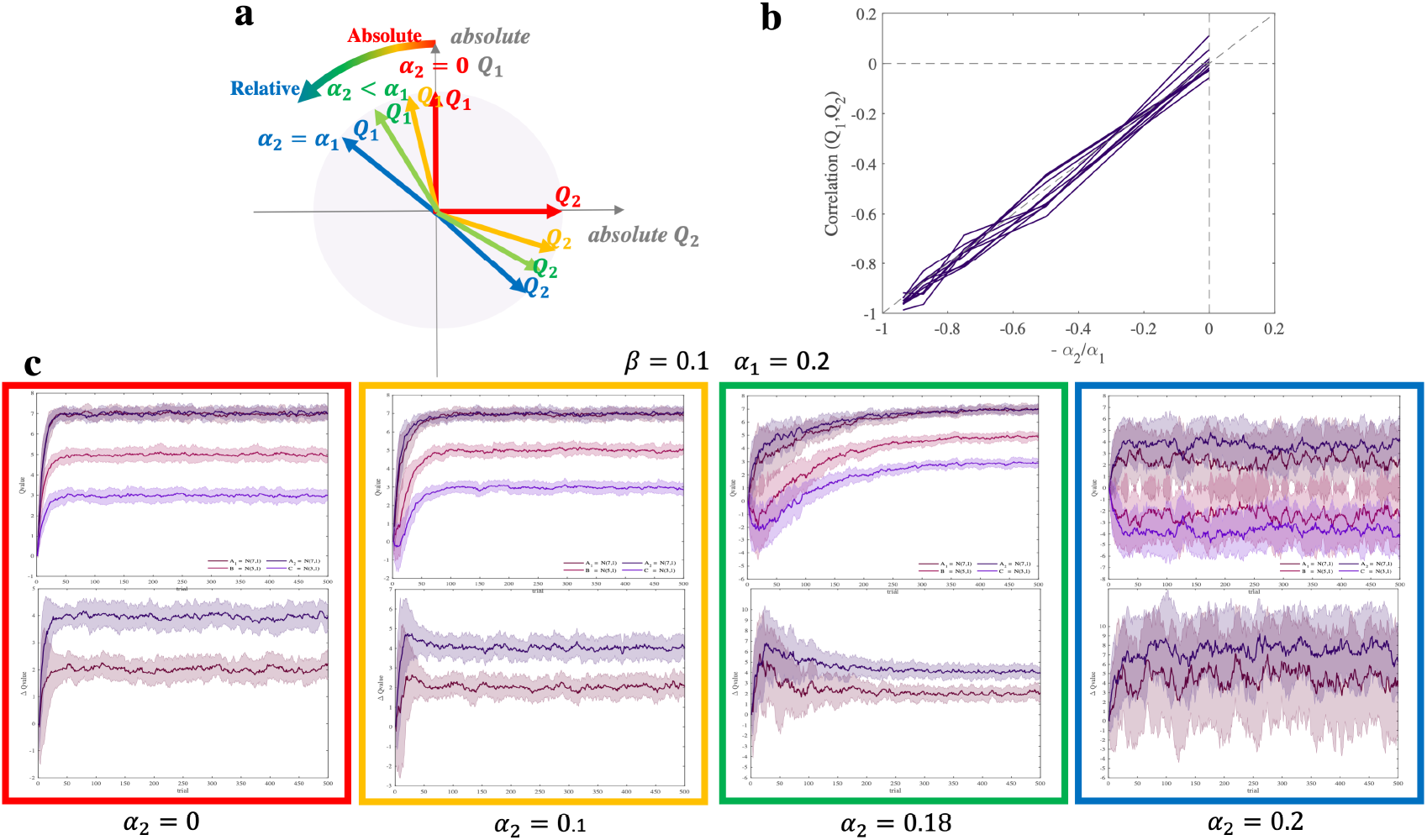
Correlation between two competing options’ values estimated by the OL model. **a.** When *α*_2_ = 0, two estimated values are equal to their absolute values and they are orthogonal. But whenever *α*_2_ gets closer to *α*_1_, the estimated values in each pair become more correlated and each of them represents a stronger combination of the two absolute values. And when *α*_2_ = *α*_1_, estimated values are approximately fully correlated (*corr* ≈ −1). **b.** The correlation between two paired options’ values as a function of −*α*_2_*/α*_1_. **c.** The difference in the estimated values of *A*_1_ and *A*_2_ (contextual bias) emerges with increasing *α*_2_. The q-values and their differences are in top and bottom parts of the figure respectively. The simulation has been done on two different pairs of options 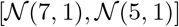, and 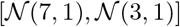, with *β* = 0.1, *α*_1_ = 0.2, and four different *α*_2_ = 0, 0.1, 0.18, 0.2.

According to this formula, the amount of correlation between *Q*_1_ and *Q*_2_ depends on the amount of the unchosen learning rate. When *α*_2_ changes from 0, where *Q*_1_ and *Q*_2_ are almost orthogonal (*corr* ≈ 0), to *α*_1_, where *Q*_1_ and *Q*_2_ are almost fully correlated *corr* ≈ −1), the encoding will change from almost fully absolute to almost fully relative (Figure 5a,b). Through simulating the experiment with typical agents of *α*_2_ = 0, 0 < *α*_2_ < *α*_1_, and *α*_2_ = *α*_1_, we showed that we will have zero, moderate and large amount of contextual effect with never, temporary and permanent contextual effect durations, respectively (Figure 5c, in the red box there is no contextual effect, in the yellow and green box there is a temporary moderate amount of contextual effect, and in the blue box there is a permanent large amount of contextual effect).

##### OL optimality

The inhibition role of the chosen prediction error on unchosen value would lead to an increase in the contrast between the competing options’ values, and it leads to an increase in the performance, especially in an environment within a reasonable noise range (Figure 6a). To illustrate the performance change in the OL model, we did a simulation with a wide range of task settings, *μ*_2_ − *μ*_1_ ∈ [1, 10], *σ* = 1, and a wide range of parameters, *α*_1_ ∈ [0, 1], *α*_2_ ∈ [0, *α*_1_], and *β* ∈ [0, 1] (for the full setting see the Methods). For the sake of simplicity, we performed all the simulation with the scaled Q-values (directly) and scaled *β* (inversely) so that *σ* was 1. By this scaling, the dynamics of the values will remain unchanged. Each of these simulations has been repeated 100 times and later averaged.

**Fig 6.**
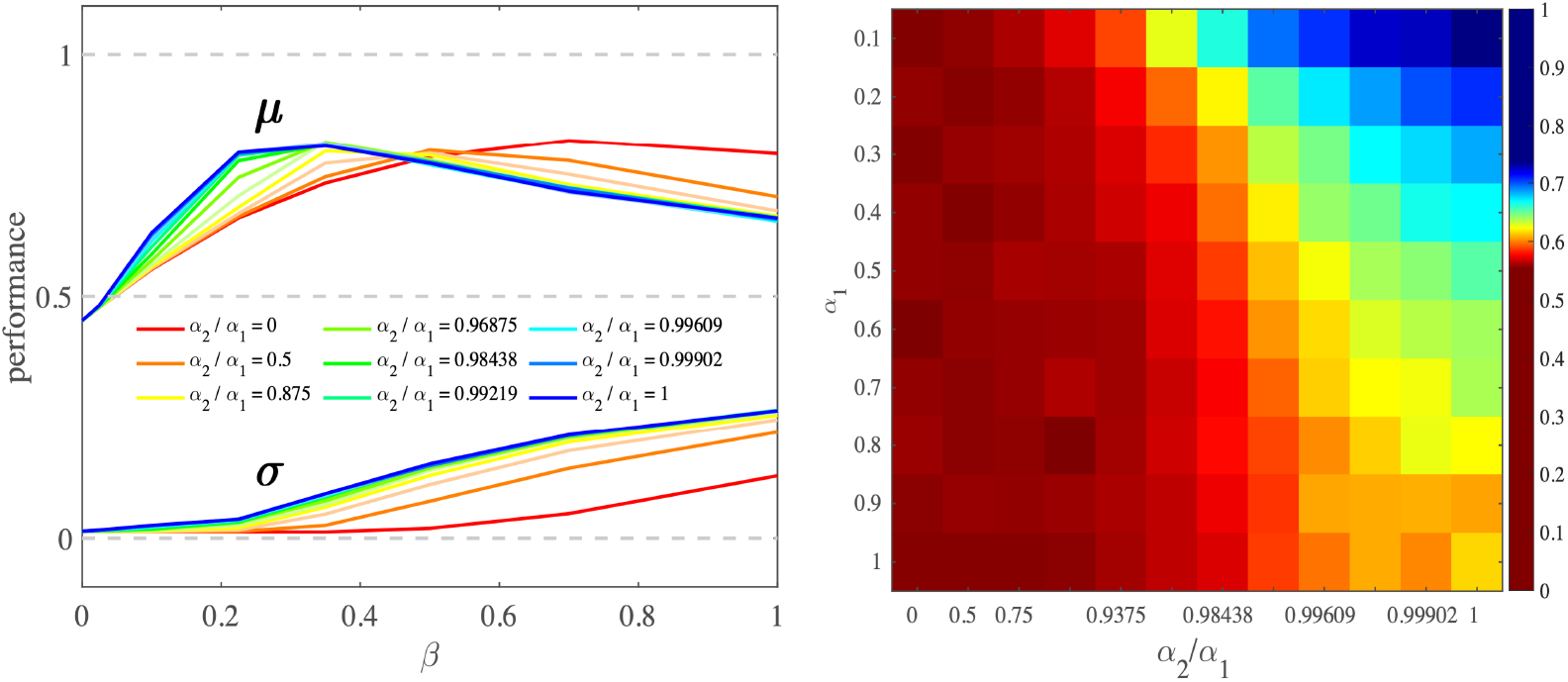
Performance comparison between OL and SQL model. **a.** As *α*_2_*/α*_1_ goes from 0 (SQL) to 1 (the OL_1_) the peak of the performance shifts to the left, where the value of *β* is smaller, and also is more reasonable. In this *β* range. For higher *α*_2_*/α*_1_ the peak of performance has been reached in higher *β* that there is high variance in behavior. The performance has been obtained by averaging performance across all task settings and different ranges of *α*_2_*/α*_1_. **b.** This heat-map shows that by increasing *α*_2_*/α*_1_, performance will increase. This result comes from the task setting 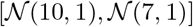, and *β* = 0.1.

Our simulation analysis in the first step showed that the OL model as a reinforcement learning model has a better performance when the difference between competing options’ values increases (Figure 1). This analysis also showed that when noise is in a reasonable range, with *α*_2_, increasing, the performance will increase as well (relative to the SQL model, Figure 6a,b), and it means by embedding the *α*_2_, inhibition mechanism in the model, we can have a more optimal learning behavior.

##### OL extension

The basic OL model introduced above, suggests the endogenous relative encoding in the Partial version. The main idea is the non-selective and diffusive behavior of dopaminergic signals on D1- and D2-SPN neurons. But in the Complete version there is another relativity inducing factor and that is to what extent factual outcomes are better or worse than the counterfactual outcomes. It has been shown that dopaminergic signals in the presence of counterfactual outcome differs from the standard prediction error, and it is the integration of reward and counterfactual prediction errors [30]. Furthermore, some studies have shown that by adding the outcome difference strategy to the learning procedure, the model can better account for the behavioral [9] or physiological [27] data. Therefore, we inserted the outcome difference component into the OL model to extend it for the Complete version (Figure 4b). It is worth mentioning that the outcome difference factor had a significant effect on the participants’ switching behavior in the Complete version and not in the Partial version.

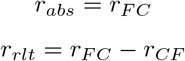

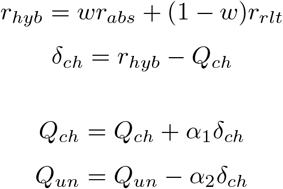

where *w* is the weight of absolute strategy. If the means of reward distributions of paired options are *μ*_1_, and *μ*_2_, and then their difference is *μ*_1_ − *μ*_2_, the means of the new reward distributions and their difference would be:

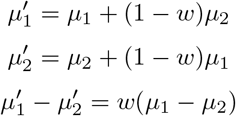

Using *r_hyb_* in the prediction error formula seems as if we have two options with two new reward distributions, in a way that that their means get closer to each other, relative to when we use *r_FC_*. Thus, this modification does not change the key OL behavior, and the extended-OL model still preserves all the above-mentioned properties. Therefore, by designing a proper prediction error, the OL model will have a good ability to be extended easily to a wide range of conditions.

#### Model comparison

##### Model fitting and model validation

In this part of the analysis, we compared the novel OL model with the related previously introduced models, in two ways: model-fitting and model-validation. We included the standard Q-learning model (SQL) as a benchmark, and the reference-point model (RP) [11], difference model [10], and hybrid model [9] as rivals in the model space. Almost all of the models had Partial and Complete versions. The OL model has two different versions, OL_1_ where the chosen and unchosen options have the same learning rates, and OL_2_ where they have different learning rates.

We did the fitting procedure for the learning phase of each subject and each model, and calculated their Bayesian exceedance probabilities. For the transfer phase, the negative log-likelihood were obtained by the likelihood that the model chooses the options that the subject has chosen in the transfer phase on its first iteration. Through model comparison, we found that the OL models especially OL_1_ (for the Partial and Complete versions), had a better fitting criterion in the learning phase and also a better prediction criterion in the transfer phase (table 2).

**Table 2.**
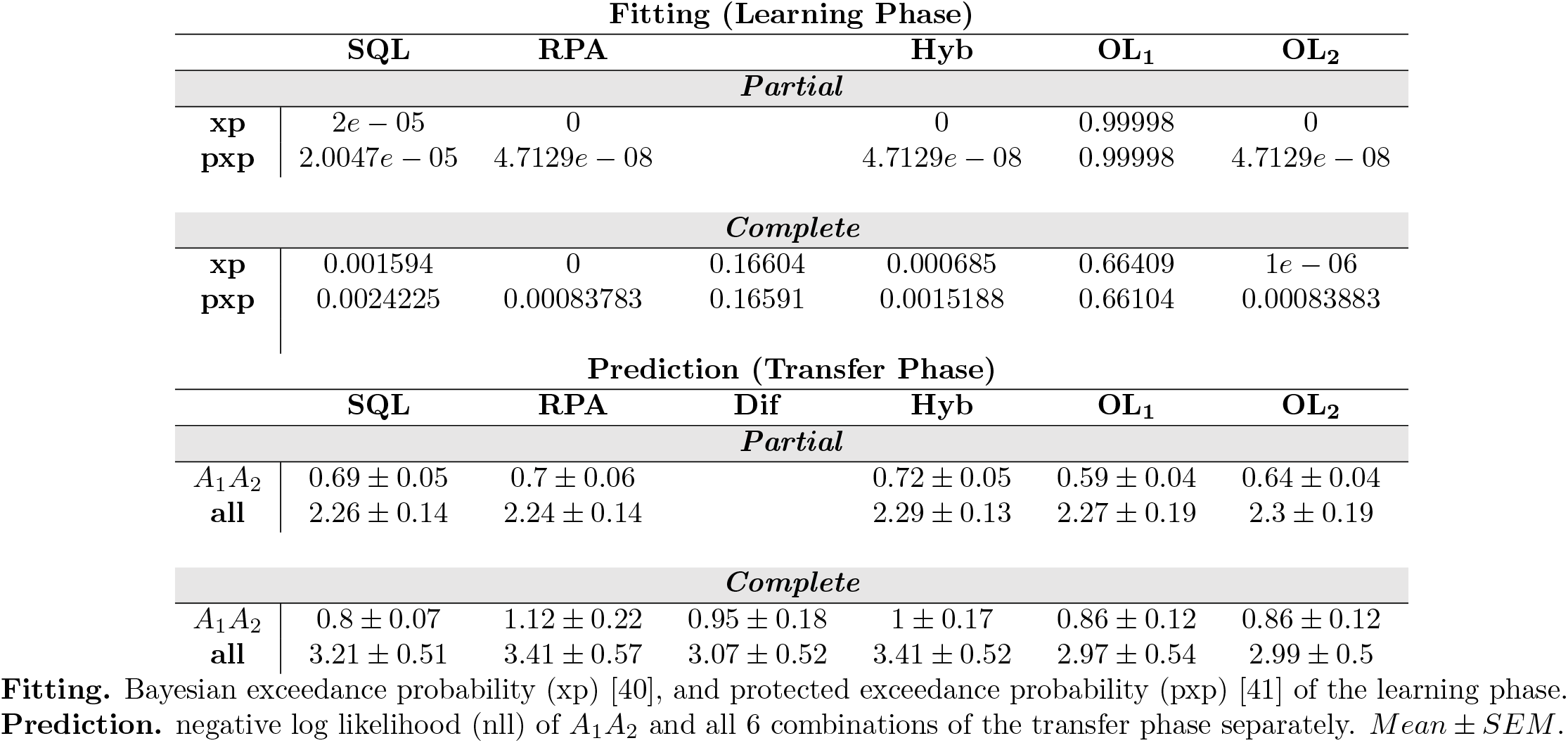
Model comparison: model fitting and model prediction.

**Table 3.** Estimated parameters.

| parameter | constraint | SQL | RPA | Dif | Hyb | OL<sub>1</sub> | OL<sub>2</sub> |
| --- | --- | --- | --- | --- | --- | --- | --- |
| *Partial* |  |  |  |  |  |  |  |
| $\beta$ | $0 \leq \beta < \inf$ | $0.07 \pm 0.03$ | $0.12 \pm 0.08$ | | $0.06 \pm 0.04$ | $0.02 \pm 0.02$ | $0.03 \pm 0.02$ |
| $\alpha_1$ | $0 \leq \alpha_1 \leq 1$ | $0.25 \pm 0.26$ | $0.26 \pm 0.27$ | | $0.37 \pm 0.29$ | $0.26 \pm 0.2$ | $0.32 \pm 0.23$ |
| $\alpha_2$ | $0 < \alpha_2 \leq \alpha_1$ | | $0.34 \pm 0.3$ | | | | $0.21 \pm 0.18$ |
| $w$ | $0 \leq w \leq 1$ | | | | $0.55 \pm 0.37$ | | |
| *Complete* |  |  |  |  |  |  |  |
| $\beta$ | $0 \leq \beta < \inf$ | $0.12 \pm 0.09$ | $0.37 \pm 0.24$ | $0.37 \pm 0.23$ | $0.2 \pm 0.15$ | $0.11 \pm 0.12$ | $0.1 \pm 0.1$ |
| $\alpha_1$ | $0 \leq \alpha_1 \leq 1$ | $0.14 \pm 0.16$ | $0.1 \pm 0.12$ | $0.09 \pm 0.08$ | $0.21 \pm 0.15$ | $0.22 \pm 0.15$ | $0.26 \pm 0.14$ |
| $\alpha_2$ | $0 < \alpha_2 \leq \alpha_1$ | | $0.11 \pm 0.13$ | | | | $0.19 \pm 0.16$ |
| $\alpha_3$ | $0 \leq \alpha_3 \leq 1$ | | $0.35 \pm 0.3$ | | | | |
| $w$ | $0 \leq w \leq 1$ | | | | $0.28 \pm 0.23$ | $0.28 \pm 0.17$ | $0.32 \pm 0.19$ |

The estimated parameters for each model. *Mean* ± *SD*.

In addition to model fitting analysis, we used model-validation analysis to test whether the OL model can generate the observed behavior. The simulation for each participant in each model was conducted by her best-fitted parameters, 100 times, and then were averaged. As expected, in the learning phase of both versions, agents’ performances were higher than 0.5 (Partial: performance = 0.6637 ± 0.0627; Complete: performance = 0.8857 ± 0.0639; Figure 3b), and consistent with the behavioral results, the performance in the learning phase of the Complete version was significantly higher than that in the Partial version (*p* = 4.4086*e* − 25, *tstat* = 15.3079, *df* = 75, one-tailed ttest). In addition to the learning phase, we also observed that the performance of the subjects was high in the transfer phase, such that participants significantly preferred the option with higher expected values (Partial: *p* = 5.4079*e* − 105, *tstat* = 31.8008,*df* = 348; Complete: *p* = 3.1818*e* − 177, *tstat* = 49.5978, *df* = 418; binomial test). We could also replicate the transfer effect (Figure 3a), in a way that agents preferred *A*_2_ over *A*_1_ in both feedback versions (Partial: *p* = 0.04096, *ratio* = 0.65714; Complete: *p* = 6.8771*e* − 05, *ratio* = 0.78571; binomial test). This simulation analysis showed that the OL model could generate all key signatures of the behavioral data (Figure 3a,b).

##### Parameter recovery

To validate our model fitting, we probed the correlation between fitted and recovered parameters. For each best-fitted parameter, we performed parameter recovery for 100 distinct simulations and then averaged it. We found strong correlations between fitted and recovered parameters, (*corr* ≥ 0.9) (Figure 7).

**Fig 7.**
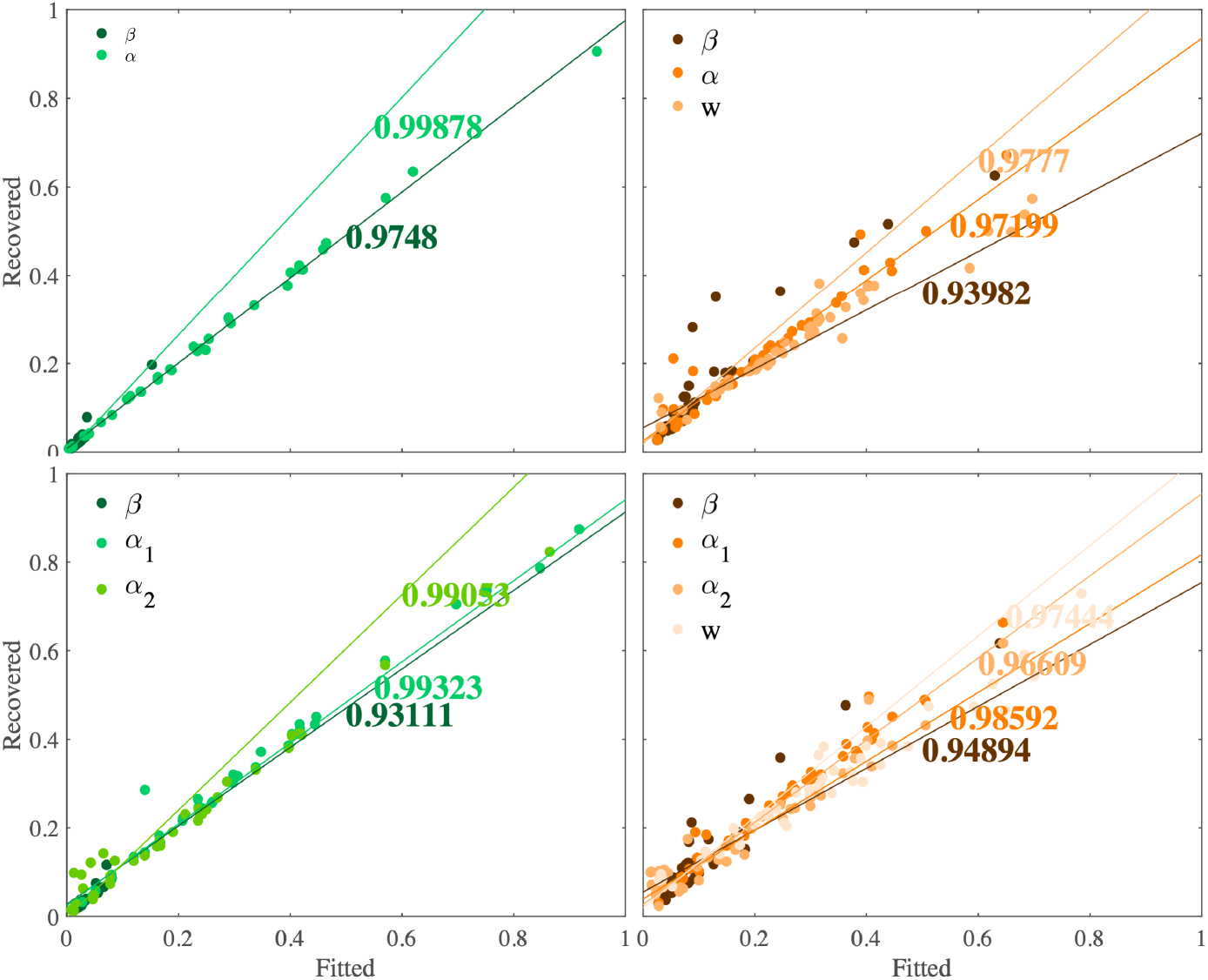
The correlations between fitted and recovered parameters. The OL_1_ (top) and OL_2_ (down) model for the Partial (left, green), and Complete (right, brown) versions. The recovery for each fitted parameter has been done 100 times and then averaged.

## Discussion

The investigations of contextual effect on value learning have mostly focused on the putative role of counterfactual components in the Complete version. In this study, we showed that counterfactual components play an important role also in the Partial version where only factual outcomes are provided, and the counterfactual component here is the effect of chosen outcome on unchosen value. Inspired by the opposing role of dopamine on competing options’ values in the striatum, we introduced a novel Opposing Learning model, in which the chosen prediction error, updates the competing options’ values in an opposing manner. Unchosen value updating with chosen prediction error will make the competing options’ values correlated to each other which leads to the emergence of the contextual effect during learning. On the other hand, due to the inhibiting role of the prediction error in unchosen values, the contrast between options’ values compared to the standard Q-learning model will increase, and this leads to higher performance in a reasonable exploration rate and more optimal learning than the standard way. This model could show the behavioral characteristics of the data and also by comparing it with the previous related models, it could better account for the data.

The majority of studies on instrumental learning paradigm used discrete rewards of 1 and 0 as gain and loss and subjects were supposed to estimate the probability of rewards for each option to maximize their payoffs [10, 11, 31]. But in the real world, we often experience continuous outcomes of our choices and are supposed to estimate their expected outcomes. Our secondary aim in this study was then to investigate the contextual effect in the paradigm with continuous reward amplitudes. We adopted previous instrumental learning tasks with novel reward designs, in which the stimuli were associated with some rewards drawn from specific normal distributions. With these complementary results, we could show that the contextual effect is not limited to probabilistic reward, but it extends to reward with amplitude.

Learning and decision making are two intermingled processes, and studying either of them cannot be separated from the other, as the recent evidence showed that some decision-making biases come from value learning that happens in a specific context [9–11]. The main neural underpinnings of these two processes are in the striatal circuitry in the subcortical part of the brain [42, 43]. A wide range of studies have shown a correspondence between the well-known reinforcement learning model, and striatum function [24, 26]. Dopamine is proposed to encode reward prediction errors [18, 44] and it reinforces the representations in the striatum [45], the region that has been proposed to encode options’ values [12–17]. The main assumption in this model is that the chosen prediction error only affects the chosen value. But it has been proposed that the underlying function in the striatum relies on the opposing role of dopamine on two segregated populations of neurons which encode the competing options’ values separately [21, 22, 25, 46]. This encouraged us to attempt to modify the standard Q-learning model to have a model more consistent with physiological evidence.

There are two direct and indirect pathways in the Basal Ganglia which have shown to have opposing roles; direct pathway promotes and indirect pathway inhibits options [24, 26]. These pathways originate from two distinct populations of neurons in the striatum, D1-SPNs, and D2-SPNs respectively, in which dopamine has an opposing influence on them, by stimulating D1-SPNs, and inhibiting D2-SPNs neurons [19, 20]. In associative learning studies, it has been shown that D1-SPNs and D2-SPNs neurons encode two opposing options’ values of competing options in a two-forced choice operant learning task [21, 22, 25, 31, 46–48], in which D1-SPNs encodes the ongoing (chosen) option and D2-SPNs encodes its competing option. Being inspired by this evidence, we introduced a novel model in which chosen-related prediction error updates the chosen and unchosen value concurrently, but in an opposing manner by updating the latter and former in an increasing and decreasing manner respectively. The OpAL model have been previously introduced by Collins et al with a similar idea [48]. The main difference between OpAL and OL models is that OpAL uses reference-point mechanism explicitly, but in the OL model without explicit using of reference-point, it emerges during learning implicitly, and without adding complexity of reference point calculations, OL model predicts the behavior in a better manner.

The OL model having two concurrent associative learning for opposing actions has a good potential to explain the recent neural evidence. Several studies in different ways have shown that stimulation of D1-SPNs increases the approach behavior and decreases the avoid behavior, and stimulation of D2-SPNs increases avoid behavior and decreases approach behavior for the ongoing action. This evidence has also been shown by increasing and decreasing the amount of dopamine in the striatum [49], stimulating and inhibiting D1-SPNs and D2-SPNs by light [50], and removing the D1-SPNs and D2-SPNs activity by ablation experiment [51]. The relative activation of these two pathways encodes the internal variable of the underlying decision-making procedure [25], that can play the role of likelihood computation in the softmax rule [25] and make a bias towards the option with higher value [52, 53]. The specificity in these two pathways is similar and the amount and pattern of their activations are anti-correlated [23, 54, 55]. Similar kinds of reported evidence in decision-making paradigms have also been reported in the learning paradigms [50, 56, 57]. The opposing synaptic plasticity in these two pathways was also reported [58]. As has been shown in the Results Section, the OL model can potentially account for this evidence.

Due to being concurrently affected by chosen-related prediction error, competing options’ values are encoded depending on each other. Indeed, this dependency appears as a correlation that is proportioned to −*α*_2_*/α*_1_. Whenever *α*_2_ gets closer to *α*_1_, their (absolute) correlation increases, such that when *α*_2_ ≈ 0, the correlation is the least (*corr* ≈ 0), and when *α*_2_ = *α*_1_ the correlation is the most (*corr* ≈ −1). This correlation is also consistent with the physiological evidence which has shown that D1-SPNs and D2-SPNs neurons in the instrumental learning tasks have opposite activity with similar strength [21, 22]. Since in this model the competing options’ values are anti-correlated, the OL estimated values depend on their paired options, and then this model generates the contextual effect. The amount of this contextual effect is proportioned to *α*_2_*/α*_1_. When *α*_2_ = 0, there is no contextual effect at all, when 0 < *α*_2_ < *α*_1_, there is a moderate amount of contextual effect that is temporary and disappears over time (but in a long run). And when *α*_2_ = *α*_1_, there is the largest contextual effect that is permanent.

We showed that the OL model compared to its counterpart, the standard Q-learning model, has an advantage of being more optimal by having higher performance. Whenever *α*_2_ gets closer to *α*_1_, the performance in the environments with a reasonable amount of noise will increase, in a way that the more relative the model is, the higher is the performance. Improvement of performance is because of boosting the contrast between the options’ values which leads to detect the superior options. Analogous to the OL model, there is also this kind of optimal behavior in the confirmation bias model. In this model, it is the asymmetric updating of positive and negative prediction errors for chosen and unchosen options’ values that boosts the contrast between options’ values [59].

It has been shown that people are not only affected by their factual rewards but also by their relative rewards that are the difference between the factual and counterfactual outcomes [27–29]. These relative outcomes are also encoded in the brain by dopamine [30, 60]. In particular, in conditions in which this comparison is available to participants, this effect is stronger and participants use the hybrid of absolute and relative strategies to learn and choose [9, 27]. In our behavioral analysis, we showed that the comparison effect is stronger in the Complete version than in the Partial one. This exogenous relativity is a different component compared to the endogenous relativity introduced by the OL model, then by inserting this factor into the model, we can expect to have better accounting for the behavioral data. As this model can be extended to any other well-defined prediction errors and preserve all its characteristics, we extend the OL model for the Complete version by inserting the outcome comparison part to it. This embedding could better explain the Complete version.

Substantial evidence demonstrates that two separate and parallel systems are involved in decision-making and learning, the Basal Ganglia and Frontal cortex, in which the Basal Ganglia plays a critical role for habitual behavior and the Frontal cortex plays a critical role in the goal directed behavior [61]. It is the weighted combination of these two systems that are involved in people’s behavior. It has been shown that several factors modulate these weights [62–72]. Different amount of contextual effect in the learning, transfer and estimation phases are in line with this hypothesis. In each phase of the task, participants have different needs. In the learning phase, to gain more rewards, they need to know how much an option is better than its competing option. We expect to see that the learning phase strategy is reflected in the transfer phase, where they are supposed to continue to choose between pairs of options. Finally, in the estimation phase, in contrast to previous phases, they need to know the exact absolute values. According to these needs, we expect to have the most BG, the least FC weights in the learning phase, the modest BG, and FC weights in the transfer phase, and the least BG, the most FC weight in the estimation phase [73].

Taken together, in this paper we could show that we are affected by the context by the fine interaction of counterfactual outcomes. In the two-option learning tasks, we learn the value of each option relative to its alternative, even when we don’t explicitly use the comparison strategy. On the other hand, although this contextual effect results in suboptimal decision-making outside the original context, it leads to an ecological advantage by gaining more rewards within the original context. Furthermore, and not surprisingly, people can access to both relative and absolute estimations of their options’ values, and to use which of them depends on their needs and conditions. Like other contextual biases and irrationalities in the human behavior, this bias seems to have an advantage for people to use. Investigating the mechanism of these irrationalities helps us find a solution in conditions where advantages change into disadvantages, and it will be more critical when they change to disorder.

## Materials and methods

### Participants

Two groups of 41 and 47 subjects have participated for the Partial and Complete versions of our task respectively. We excluded 6 subjects from the Partial version and 5 subjects from the Complete version (2 and 3 subjects because they didn’t learn the associations, and 4 and 2 subjects because their expected rewards for *A*_1_, and *A*_2_ were more than one, in Partial and Complete versions respectively, see below). After exclusion *N* = 35 subjects (age: 26 ± 6 (*mean* ± *SD*), female: *n* = 16) and *N* = 42 subjects (age: 23 ± 5 (*mean* ± *SD*), female: *n* = 12) remained for analysis in the Partial and Complete versions respectively. They were received their monetary rewards after they completed the task, according to their performances. They were all healthy volunteers that gave a written consent before starting the task. The study was approved by the local ethics committee.

### Behavioral task

Two different cohorts of participants performed two different versions of instrumental learning tasks, which were adopted from previous studies [9–11]. The main structure of these two tasks was almost the same and included two consecutive phases of learning and post-learning transfer. The only difference was in the way feedbacks were provided to the subjects. In the Partial version, only the factual outcomes for chosen option were provided to the subjects, and in the Complete version, both the factual and counterfactual outcomes for chosen and unchosen options, respectively, were provided. Before the main task, subjects performed a short training session (20 trials) to be familiarized with the learning phase. The stimuli and the reward statistics of the training session were different from those of the main session. The stimuli were selected from the Japanese Hiragana alphabet.

The learning phase was made up of one session in which, in each trial two stimuli were presented on the screen, and participants were instructed to choose the option with higher expected rewards. This instrumental learning paradigm made participants to learn gradually by trial and error to choose the most advantageous option in each trial. The cues were shown to the subjects from two pairs of stimuli {*A*_1_*B, A*_2_*C*}, which means in each pair each stimulus was always presented with a similar stimulus. Each pair thus established a fixed context. These two contexts were pseudo-randomly interleaved across trials. The rewards of *A*_1_ and *A*_2_ stimuli were drawn from the same normal distribution of 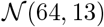 and the rewards of *B* and *C* stimuli were drawn from a different normal distributions of 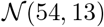 and 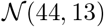, respectively. To control some confounding factors, rewards samples were drawn from the truncated distribution, which was in the [*μ* − 3*σ, μ* + 3*σ*] ([0, 100]) interval. The parameters of the distributions were unknown to the subjects, and they were supposed to learn them. Although the reward statistics of *A*_1_ and *A*_2_ were the same, the images associated with them were different to conceal the task structure from the subjects.

The side of each stimulus on the screen, whether the right of the fixation point or the left, was also pseudo-randomized during the task, such that for the total number of trials for each context, in half of the trials a particular stimulus was presented on the right and in the other half, on the left. The subjects were asked to select their choices within a 4000 ms, otherwise, they missed that trial’s reward, and the ‘No Response’ message was shown on the screen. Within each trial, the subjects chose their choice by pressing the left and right arrow keys for the left and right options respectively. Following the choice, the chosen option was surrounded with a blue square and the related outcomes were presented simultaneously on the screen. In the Partial version, the factual outcome was shown below the chosen option for 500 ms and in the Complete version, both the factual and counterfactual outcomes were shown below the chosen and unchosen options respectively for 1000 ms. In the Complete version, the information that subjects should process was two times the Partial version and in our pilot study, we found that having only 500 ms for observing the outcomes was not sufficient to process two continuous outcomes and so decreased the subjects’ performance compared to the Partial version, therefore we doubled this time to 1000 ms. The next trial started after 1000 ms fixation screen. Each context was presented to the subjects at least in 50 trials and then two contexts consist of, at least 100 trials. After at least 100 trials, the task continued for each subject until the experienced mean of *A*_1_ became almost equal to the experienced mean of *A*_2_, (their difference became less than 1). If this condition was not met up in the 300th trial then the learning phase was stopped and this subject was excluded from the data. By this design, the number of trials always falls into the range of [100,300] and this number might be different for each subject.

Seamlessly after the learning phase, participants entered the post-learning transfer phase. They were not aware of the transfer phase until they completed the learning phase, in order not to use any memorizing strategy in the learning phase. In the transfer phase, all possible binary combinations of the stimuli (6 combinations) were presented to the participants and they were asked to choose the option with higher expected rewards. They were told that they will not only see the previously paired options in the learning phase but even the binary options which weren’t paired in the preceding phase. Each combination was presented four times, giving a total 6 * 4 = 24 trials that were presented in a pseudo-randomized order. This phase in contrast to the learning phase was self-paced (they were not force to choose in a limited time) and also no feedback was provided to the subjects, in order not to interfere with their learned values [9–11, 31, 32]. Following each choice, they had to report the confidence of their choice by using a scaled bar from 0 to 100 in which the leftmost side of the axis shows complete uncertain and the rightmost side shows complete certain. The confidence part was done by the mouse. After the transfer phase, subjects completed the estimation phase. In the estimation phase, stimuli were presented to the subjects one by one and they were asked to estimate its mean of rewards, using a scaled bar from 0 to 100. Each stimulus was repeated four times giving a total of 4 * 4 = 16 trials which were presented pseudo-randomly. These trials were also self-paced and no feedback were provided to the subjects. The subjects were told their payoffs are based on the sum of rewards they would gain during the learning task. In the Complete version, subjects were notified that their total rewards are only based on the rewards of their choices. Although they were not paid in the transfer phase, they were encouraged to do as best as they can to answer correctly as if they would be paid. At the end of the task, their total rewards were shown on the screen.

### Computational models

#### The Standard Q-Learning (SQL) Model

It is a common approach to compare the context-dependent learning models with the standard Q-learning model as a benchmark that plays the role of absolute learning model. In this model, the value of each option is only related to its own observed outcomes and not to other alternative outcomes.

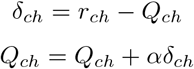

In the simplest form, it is only the chosen option which is updated following its outcomes observation, while in its extended form the unchosen options are also updated, but again with their own observed outcomes:

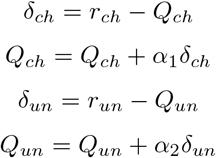

Which their learning rates can be the same or different (*α*_1_ = *α*_2_ or *α*_1_ ≠ *α*_2_).

#### The Reference-Point (RP) Model

The idea of the reference-point (RP) model comes from the reference point phenomenon which is reported by behavioral and economic studies [74, 75]. According to this model, there is a distinct reference-point for each context that is obtained by its expected outcomes. Then, the relative outcome of each option is calculated in comparison to this reference-point. We implemented several forms of RP models considering the several forms of context reward [11]. The RPD, RPA, and RPM, when the contextual rewards, *r_x_*, are considered to be direct *r_ch_*, average of (*r_ch_* + *Q_un_*)/2, and max(*r_ch_, Q_un_*) respectively in the Partial version, and *r_ch_*, (*r_ch_* + *r_un_*)/2, and max(*r_ch_, r_un_*) in the Complete version.

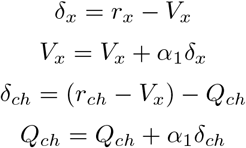

where *V_x_* is the value of the context, and *Q_ch_* is the value of the chosen option. For the Complete version, we also update the unchosen options as below,

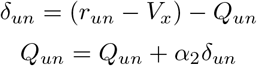

In the Complete version, we used different versions for RP. One which only updates the chosen value, and one which updates both options with the same and different learning rates.

#### The Difference (Dif) Model

Learning in a specific context in which a participant is supposed to maximize her rewards needs using a strategy in order to find a better option as soon as possible. The difference model is one of the models which gives a fast detection of the advantageous option by learning the relative value. In this model, the participants learn how much the superior option is better than the inferior one [10].

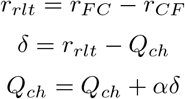

This model was only applied for the Complete version.

#### The Hybrid (Hyb) Model

It has been shown that people are not fully absolute or fully relative learners, rather they are hybrid learners in which their behaviors depend on how much they weigh either of these strategies [9].

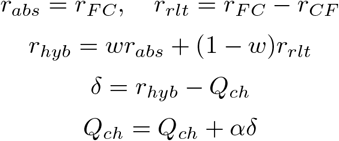

For the Partial version, we used the *Q_un_* instead of *r_CF_*.

#### The Opposing Learning (OL) Model

The OL model has been inspired by the opposing role of dopamine as prediction error on the chosen and unchosen options. In this model, the unchosen option is updated simultaneously with the chosen option and proportional to the chosen prediction error, but in an opposite manner.

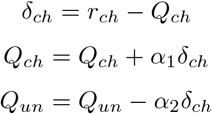

In this model, the *α*_2_ parameter controls the amount of contextual effect on the value learning procedures. For the Complete version, this model was extended to a version in which the counterfactual outcomes were considered in a hybrid manner.

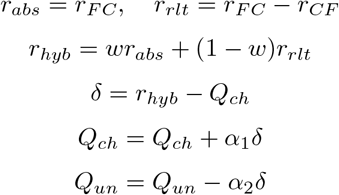

### Pure simulation procedure

The OL behavior has been examined in a wide range of task and parameter settings. Without loss of generality, we did the simulation with normalized settings such that we had *σ* = 1 in reward distributions. As an example, the normalized version of the setting of task 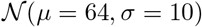, parameters of *β* = 0.01, and any *α*_1_, *α*_2_, changes to its normalized version of 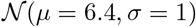 (divide by 10), and parameters of *β* = 0.1 (multiply by 10), and the same *α*_1_, *α*_2_. The tasks settings covered 10 different pairs of options in which their relative values were covered {1, 2, …, 10} ([*μ*_1_, *μ*_2_] ∈ {[10, 9], [10, 8], …, [10, 0]}, and *σ* = 1). The parameters settings covered a wide range of *β*: {0, 0.025, 0.05, 0.075, 0.1, 0.1025, …, 0.4} ∪ {0.5, 0.6, …, 1}, *α*_1_: {0.1, 0.2, …, 1}, and *α*_2_*/α*_1_: {0, 0.5, 0.75, 0.875, 0.93, 0.96, 0.980.992, 0.996, 0.998, 0.999, 1}.

### Fitting and simulation procedure

The data fitting was implemented by *fmincon* function of Matlab software (the MathWorks Inc., Natick, MA). The fittings have been done with several initial points to have higher probability in order to find a global optimum, rather than getting stuck on a local optimum. For obtaining the exceedance probabilities (xp) [40], and protected exceedance probabilities(pxp) [41] for the model-comparison part, and estimating parameters, we optimized maximum a posteriori (MAP) using weakly informative priors of *β*(1.2, 1.2) for each parameter. It is worth noting that the range of options’ values is in scale of 100, and so range of the *β* parameter will be in scale of much less than one, thus, the *β*(1.2, 1.2) would be a proper prior in model fitting. The exceedance probability and protected exceedance probability have calculated based on [40, 41]. The simulation for each subject was done on its best fitted parameters for 100 repetitions, and then the representative behavior of this agent was obtained by averaging across its repetitions.

## Additional information

Supplementary information is available for this paper.

## Supporting information

**S1 Table.** Model comparison with BIC.

| *Partial* |  |  |  |  |
| --- | --- | --- | --- | --- |
|  | nll | BIC |  |  |
| | learning | learning | learning + transfer($A_1A_2$)) | learning + transfer(all)) |
| **SQL** | $88.18 \pm 5.49$ | $186.82 \pm 11.05$ | $188.21 \pm 11.05$ | $191.37 \pm 11.03$ |
| **RPD** | $87.17 \pm 5.49$ | $190.04 \pm 11.11$ | $191.48 \pm 11.1$ | $194.69 \pm 11.1$ |
| **RPA** | $87.69 \pm 5.47$ | $191.07 \pm 11.06$ | $192.51 \pm 11.07$ | $195.62 \pm 11.06$ |
| **RPM** | $87.18 \pm 5.49$ | $190.05 \pm 11.11$ | $191.48 \pm 11.1$ | $194.69 \pm 11.1$ |
| **Hyb** | $86.68 \pm 5.48$ | $189.05 \pm 11.07$ | $190.5 \pm 11.08$ | $193.7 \pm 11.06$ |
| **OL<sub>1</sub>** | $84.7 \pm 5.49$ | $179.86 \pm 11.06$ | $181.05 \pm 11.06$ | $184.45 \pm 10.99$ |
| **OL<sub>2</sub>** | $83.66 \pm 5.37$ | $183.01 \pm 10.86$ | $184.3 \pm 10.86$ | $187.73 \pm 10.8$ |
| *Complete* |  |  |  |  |
|  | nll | BIC |  |  |
| | learning | learning | learning + transfer($A_1A_2$)) | learning + transfer(all)) |
| **SQL** | $54.34 \pm 4.98$ | $119.24 \pm 9.99$ | $120.84 \pm 10.01$ | $125.72 \pm 9.95$ |
| **QL<sub>21</sub>** | $51.71 \pm 4.99$ | $113.98 \pm 10.01$ | $116.19 \pm 9.98$ | $121.69 \pm 9.95$ |
| **QL<sub>22</sub>** | $50.11 \pm 5.01$ | $116.05 \pm 10.08$ | $118.4 \pm 10.03$ | $124.06 \pm 10$ |
| **RPA<sub>1</sub>** | $51.71 \pm 4.99$ | $119.25 \pm 10.03$ | $121.18 \pm 9.98$ | $125.53 \pm 9.9$ |
| **RPA<sub>2</sub>** | $48.45 \pm 4.99$ | $118 \pm 10.05$ | $120.26 \pm 9.98$ | $124.96 \pm 9.87$ |
| **RPM<sub>1</sub>** | $51.71 \pm 4.99$ | $119.25 \pm 10.03$ | $120.84 \pm 10$ | $125.55 \pm 9.92$ |
| **RPM<sub>2</sub>** | $47.81 \pm 5$ | $116.73 \pm 10.07$ | $118.59 \pm 10.03$ | $124.54 \pm 9.86$ |
| **Dif** | $51.71 \pm 4.99$ | $113.98 \pm 10.01$ | $115.89 \pm 9.95$ | $120.18 \pm 9.87$ |
| **Hyb** | $48.94 \pm 5$ | $113.72 \pm 10.05$ | $115.73 \pm 10.02$ | $120.63 \pm 9.91$ |
| **OL<sub>1</sub>** | $47.98 \pm 5$ | $111.79 \pm 10.05$ | $113.52 \pm 10.02$ | $117.83 \pm 9.93$ |
| **OL<sub>2</sub>** | $47.53 \pm 4.96$ | $116.17 \pm 9.99$ | $117.9 \pm 9.97$ | $122.27 \pm 9.88$ |

BIC of three different parts, learning phase, learning and ( $A_1A_2$ ) of the transfer phase, and learning and all 6 combinations of the transfer phase for model-space.

**S1 Fig.**
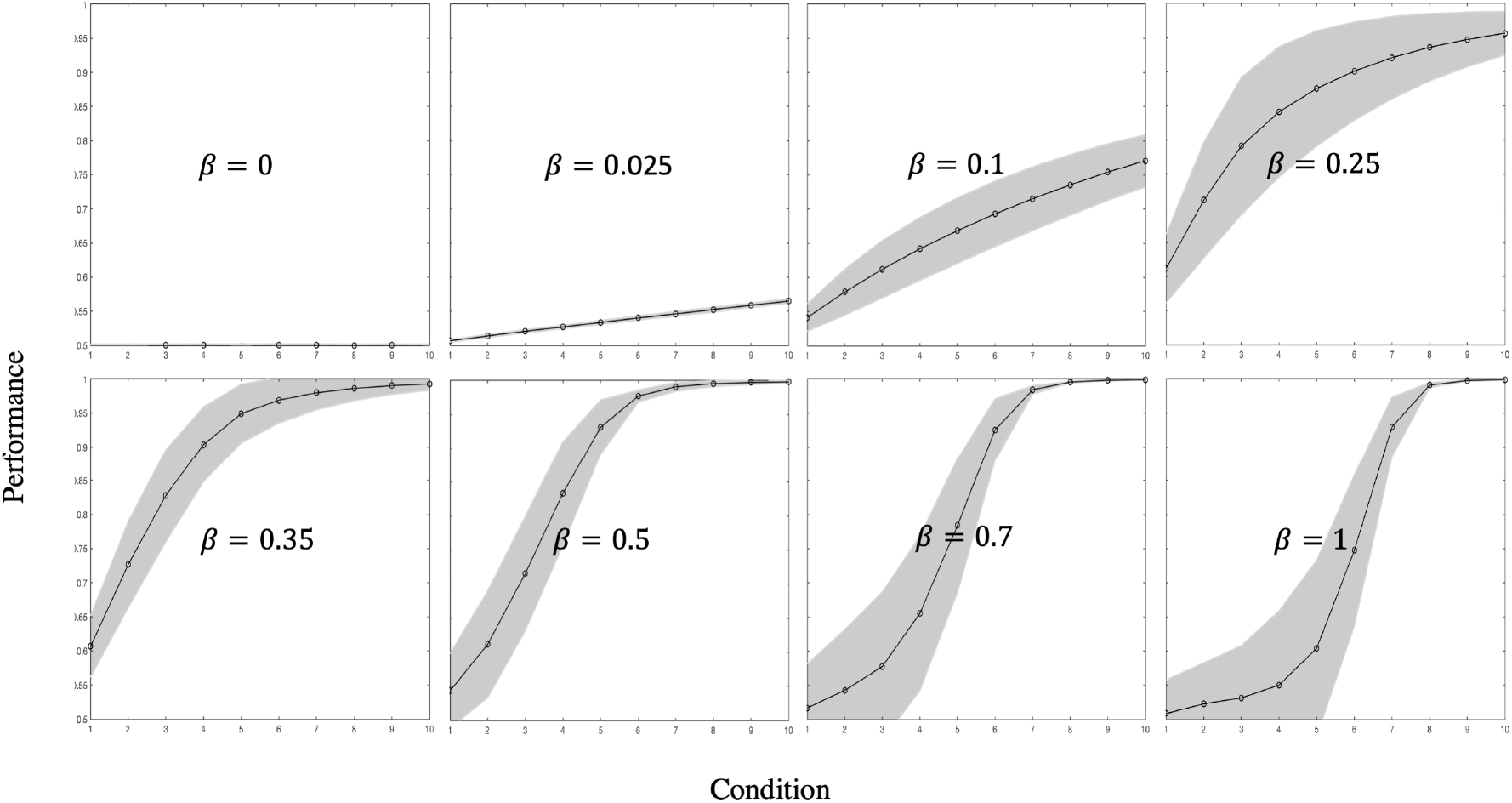
An OL agent has higher performance when the distance between options values are higher. The function of the performance changes with *β* as a variable. The conditions were covered 10 different pairs of options in which their relative values were covered {1, 2, …, 10} ([*μ*_1_, *μ*_2_] ∈ {[10, 9], [10, 8], …, [10, 0]}, and *δ* = 1). Performances were obtained with averaging across different *α*_1_ and *α*_2_.

